# Dysregulated Oscillatory Connectivity in the Visual System in Autism Spectrum Disorder

**DOI:** 10.1101/440586

**Authors:** R.A. Seymour, G. Rippon, G. Gooding-Williams, J.M. Schoffelen, K. Kessler

## Abstract

Autism Spectrum Disorder is increasingly associated with atypical perceptual and sensory symptoms. Here we explore the hypothesis that aberrant sensory processing in Autism Spectrum Disorder could be linked to atypical intra- (local) and inter-regional (global) brain connectivity. To elucidate oscillatory dynamics and connectivity in the visual domain we used magnetoencephalography and a simple visual grating paradigm with a group of 18 adolescent autistic participants and 18 typically developing controls. Both groups showed similar increases in gamma (40-80Hz) and decreases in alpha (8-13Hz) frequency power in occipital cortex. However, systematic group differences emerged when analysing intra- and inter-regional connectivity in detail. Firstly, directed connectivity was estimated using non-parametric Granger causality between visual areas V1 and V4. Feedforward V1-to-V4 connectivity, mediated by gamma oscillations, was equivalent between Autism Spectrum Disorder and control groups, but importantly, feedback V4-to-V1 connectivity, mediated by alpha (8-13Hz) oscillations, was significantly reduced in the Autism Spectrum Disorder group. This reduction was positively correlated with autistic quotient scores, consistent with an atypical visual hierarchy in autism, characterised by reduced top-down modulation of visual input via alpha-band oscillations. Secondly, at the local level in V1, coupling of alpha-phase to gamma amplitude (alpha-gamma phase amplitude coupling, PAC) was reduced in the Autism Spectrum Disorder group. This implies dysregulated local visual processing, with gamma oscillations decoupled from patterns of wider alpha-band phase synchrony (i.e. reduced PAC), possibly due to an excitation-inhibition imbalance. More generally, these results are in agreement with predictive coding accounts of neurotypical perception and indicate that visual processes in autism are less modulated by contextual feedback information.

## Introduction

Autism Spectrum Disorder (ASD) is a life-long neurodevelopmental condition, characterised by impairments in social interaction and communication, and the presence of repetitive patterns of behaviours, interests or activities (APA 2013). Although these features remain the primary diagnostic markers of ASD, the presence of sensory symptoms have recently been given a more central role, consistent with findings of autism-related individual differences in visual perception, see Robertson and Baron-Cohen (2017). Additionally, over 90% of ASD individuals experience hyper- and/or hypo-sensitive responses to certain stimuli, which can result in sensory overload (Leekam *et al*., 2007). Differences in central coherence, local/global biases and predictive coding have all been proposed as possible mechanisms for these sensory symptoms (Happé, 2005; Mottron *et al*., 2006; Pellicano and Burr, 2012). An understanding of the neural circuits involved will prove fruitful for ASD research, and could even provide early diagnostic markers (Roberts *et al*., 2010; Kessler *et al*., 2016).

Dysregulated neural oscillations – rhythmical changes in neural activity – are a promising neural correlate of atypical perceptual processes in autism (reviewed in Kessler *et al*., 2016; Simon and Wallace, 2016). In particular, there has been increasing interest in characterising patterns of atypical high-frequency gamma-band activity (GBA, >40Hz) in ASD. Gamma oscillations play an important role in ‘temporal binding’ during sensory processing – the formation of a coherent percept essential for accurate information processing. GBA has therefore been proposed as a useful candidate frequency for studying temporal binding deficits in ASD alongside sensory symptoms more generally (Brock *et al*., 2002). At the cellular level, gamma oscillations are generated through the coordinated interactions between excitatory and inhibitory populations of neurons (Buzsáki & Wang, 2012). Therefore, findings of abnormal GBA in ASD would link with theories of an excitation-inhibition imbalance and atypical connectivity in ASD (Rippon et al; 2007).

As hypothesized, early studies of visual processing in ASD reported atypical, localised GBA responses to ‘task relevant’ stimuli as well as non-discriminant GBA increases to ‘task irrelevant’ stimuli (Grice *et al*., 2001; Brown *et al*., 2005). This was interpreted as an inability to synchronise visual responses at gamma frequencies, and bind perceptual processes into a coherent whole (Brock *et al*., 2002). A later study by Sun et al (2012), using MEG, reported reduced gamma coherence in ASD participants viewing Mooney faces. Reduced gamma coherence in the visual cortex was also reported by Peiker et al (2015a), who utilised a paradigm requiring the identification of moving objects presented through a narrow slit, necessitating the integration of perceptual information across time. However, another study by the same group, reported *greater* modulation of total gamma power in response to visual motion intensity for ASD participants (Peiker, Schneider, et al., 2015b). Furthermore an MEG study using a higher-level visuospatial reasoning task in young children, reported increased patterns of gamma-band coherence between occipital and frontal sensors in ASD (Takesaki et al., 2016). Whilst there is clear evidence of anomalous GBA during visual processing in ASD, the exact nature of these anomalies remains unclear: both increases and decreases in gamma-band power and coherence have been reported (reviewed in Kessler et al, 2016; Simon and Wallace, 2016). We suggest that shifting the focus from within-band oscillatory power towards considering oscillation-mediated functional connectivity and between-band oscillatory relationships could help with understanding oscillopathies in ASD in more detail (Kessler *et al*., 2016; Simon and Wallace, 2016).

Functional connectivity has been proposed as a unifying framework for autism, with the predominant theory emerging from fMRI data being a global reduction but local increase in connectivity (Courchesne and Pierce, 2005; Hughes, 2007). Recent M/EEG research has supported the first of these claims with reductions in global connectivity during set-shifting, slit-viewing, face processing and whole-brain resting state studies (Doesburg *et al*., 2013; Khan *et al*., 2013; Kitzbichler *et al*., 2015; Peiker *et al*., 2015). These reductions in connectivity are generally tied to feedback processes, located within the frontal lobes, and mediated by oscillations in theta (3-6Hz), alpha (8-13Hz) and beta-bands (13-30Hz). A recent study showed that during somatosensory stimulation, feedforward connectivity from primary to secondary somatosensory cortex is *increased* in ASD (Khan *et al*., 2015). This suggests that feedforward pathways in the autistic brain may be over-compensating for the lack of feedback connectivity. At the local level, M/EEG studies (e.g. Khan *et al.*, 2013; reviewed in Kessler et al., 2016) have not consistently supported the local increase in connectivity reported using fMRI (e.g. Keown *et al*., 2013). While some studies have identified patterns of GBA consistent with localised hyper-reactivity (Orekhova et al, 2007; Cornew et al, 2012), other studies report results consistent with reduced connectivity at the local as well as the global level (Khan et al, 2013). One key issue to be considered is the validity of the spectral measures of connectivity being used, as inferences based on power measures alone can be inconsistent with more complex measures of coherence/phase-locking (Port et al, 2015) or of cross-frequency coupling (Canolty and Knight, 2010).

An emerging biologically-relevant proxy for local connectivity is the coupling of oscillations from different frequency-bands, termed cross-frequency coupling (Canolty and Knight, 2010; Seymour *et al*., 2017). In particular, phase-amplitude coupling (PAC) has been proposed to act as a mechanism for the dynamic co-ordination of brain activity over multiple spatial scales, with the amplitude of high-frequency activity within local ensembles coupled to large-scale patterns of low-frequency phase synchrony (Bonnefond *et al*., 2017). Alpha-gamma PAC is also closely tied to the balance between excitatory and inhibitory (E-I) populations of neurons (Mejias *et al*., 2016), which is affected in autism (Rubenstein and Merzenich, 2003). One previous study has reported dysregulated alpha-gamma PAC in the fusiform face area during emotional face processing in autistic adolescents (Khan *et al*., 2013). Local PAC was also related to patterns of global alpha hypoconnectivity in autism, suggesting that local and global connectivity are concurrently affected. Altogether, oscillation-based functional connectivity in autism is characterised by local dysregulation and global hypoconnectivity (Kessler *et al*., 2016).

Within the context of visual processing, this view leads to several hypotheses, outlined in Kessler *et al*., (2016). Electrocorticography (ECoG) recordings in macaques and MEG in humans suggest that visual oscillations in different frequency bands have distinct cortical communication profiles. Gamma-band oscillations pass information up the visual hierarchy, in a feedforward manner, whereas alpha and beta-band oscillations mediate feedback connectivity, down the cortical hierarchy (Bastos *et al*., 2015b; Michalareas *et al*., 2016). Long-range alpha/beta connectivity has also been linked with top-down attentional processes during visual perception via the regulation of local gamma oscillations (Klimesch, 2012; Richter *et al*., 2017) and of local alpha-gamma PAC (Chacko *et al*., 2018). Hypothesising that autism is associated with alterations in directed functional connectivity (Khan *et al*., 2015), we predict reduced feedback connectivity within the visual system, mediated by oscillations in the alpha band, but potentially increased feedforward connectivity in the gamma band (Kessler *et al*., 2016). At the local level, neurotypical visual processing is accompanied by increases in alpha-gamma PAC, thought to arise through the E-I coupling between infragranular and supragranular layers of visual cortex (Mejias *et al*., 2016; Bonaiuto *et al*., 2018). Given an E-I imbalance in autism and reported local dysregulation of cortical activity, we hypothesise reduced alpha-gamma PAC within primary visual cortex in ASD participants (Khan *et al*., 2013; Kessler *et al*., 2016). Finally, if top-down alpha connectivity has a modulatory effect on bottom-up processing, then local alpha oscillations and alpha-gamma PAC, e.g. in V1, could reveal a systematic relationship with top-down alpha connectivity, e.g. from V4 (Khan *et al*., 2013). This may present itself differently between groups, with a more variable relationship between feedback connectivity and local PAC in the ASD group (Dinstein *et al*., 2012).

We tested these hypotheses using MEG, which combines excellent temporal resolution with sophisticated source localisation techniques (Van Veen *et al*., 1997; Hillebrand and Barnes, 2005). A group of 18 adolescent ASD participants and 18 typically developing controls performed an engaging visual task, to induce alpha and gamma oscillations. We characterised changes in power and connectivity between visual areas V1 and V4: two regions with strong hierarchical connectivity (Bastos *et al*., 2015; Michalareas *et al*., 2016). Additionally, we quantified local alpha-gamma PAC for V1 (Cohen, 2008; Özkurt and Schnitzler, 2011; Seymour *et al*., 2017).

## Methods and Materials

### Participants

Data were collected from 18 participants diagnosed with ASD and 18 age-matched typically developing controls, see Table 1. ASD participants had a confirmed clinical diagnosis of ASD or Asperger’s syndrome from a paediatric psychiatrist. Participants were excluded if they were taking psychiatric medication or reported epileptic symptoms. Control participants were excluded if a sibling or parent was diagnosed with ASD. Data from a further 9 participants were excluded, see *Supplementary Information*.

**Table 1:** Participant demographic and behavioural data. * = behavioural scores significantly greater in ASD>control group, t-test, p<.05.

|  | <u>N</u> | <u>Age</u> | <u>Male/Female</u> | <u>Autism</u><br><u>Quotient</u><br><u>(Adult)</u> | <u>Raven</u><br><u>Matrices</u><br><u>Score</u> | <u>Glasgow</u><br><u>Sensory</u><br><u>Score</u> | <u>Mind</u><br><u>in</u><br><u>the</u><br><u>Eyes</u><br><u>Score</u> |
| --- | --- | --- | --- | --- | --- | --- | --- |
| **ASD** | 18 | 16.67 | 14 male; 4<br>female | 32.6* | 43.8 | 65.3* | 21.8 |
| **Control** | 18 | 16.89 | 15 male; 3<br>female | 10.9 | 48.7 | 38.7 | 25.4 |

### Experimental Procedures

Experimental procedures complied with the Declaration of Helsinki and were approved by Aston University, ethics committee. Participants and a parent/guardian gave written informed consent.

### Behavioural Assessments

General non-verbal intelligence was assessed using the Raven’s Matrices Task (Raven and Court, 1998). The severity of autistic traits was assessed using the Autism Quotient (AQ; Baron-Cohen *et al.*, 2001a) and sensory traits using the Glasgow Sensory Questionnaire (GSQ; Robertson and Simmons, 2013). All 18 ASD participants, and 15 out of 18 control participants, completed the questionnaires. AQ and GSQ scores were significantly higher in the ASD group (Table 1). Participants also completed the Mind in the Eyes test (Baron-Cohen *et al*., 2001b), however, there were no group differences. The Mind in the Eyes test has been criticised for measuring emotion recognition rather than an autism-specific deficit in mental state attribution (Oakley *et al*., 2016), and therefore these scores were not analysed further.

### Paradigm

Whilst undergoing MEG, participants performed a sensory task (Figure 1A), designed to elicit gamma-band oscillations. Each trial started with a randomised fixation period (1.5, 2.5 or 3.5s), followed by the presentation of a visual grating or auditory binaural click train stimulus; however only visual data will be analysed in this article. The visual grating had a spatial frequency of 2 cycles/degree and was presented for 1.5s. To promote task engagement, cartoon pictures of aliens or astronauts were presented after the visual grating, for 0.5s but did not form part of the MEG analysis. Participants were instructed to respond to the appearance of an alien picture using a response pad (maximum response period of 1.5s). The accuracy of the response was conveyed through audio-visual feedback, followed by a 0.5s fixation period. MEG recordings lasted 12-13 minutes and included 64 trials with visual grating stimuli. Accuracy rates were above 95% for all participants.

**Figure 1:**
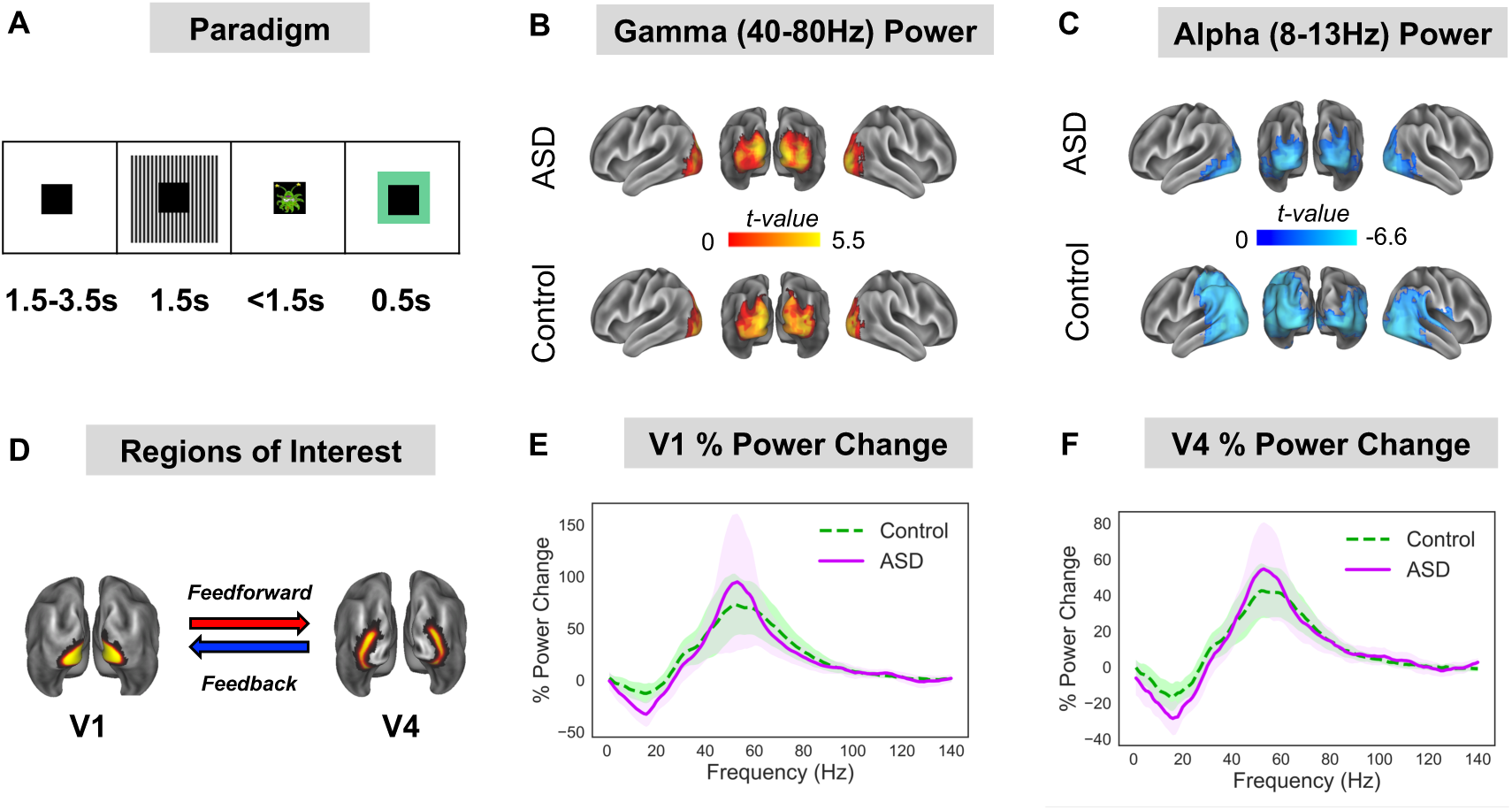
(A) Participants performed a visual task, consisting of 1.5-3.5 baseline period followed by 1.5s presentation of a visual grating. After the grating, participants were presented with a cartoon alien or astronaut picture and instructed to only respond when an alien was presented (response time up to 1.5s). The alien/astronaut stimuli were to maintain attention and do not form part of the analysis. (B-C) The change in oscillatory power between grating and baseline periods was localised on a cortical mesh and masked to show only statistically significant (p<.05, corrected) stimulus induced increases in gamma (40-80Hz) and decreases in alpha (8-13Hz) power. There were no statistically significant differences in relative gamma or alpha power between groups (see Supplementary Figure S1 for a whole-brain comparison). (D) Regions of interest in V1 and V4 were defined using HCP-MMP 1.0 atlas (43). (E-F) The change in power between grating and baseline periods was calculated for V1 and V4 from 1-140Hz. Results show characteristic reductions in alpha/beta power and increases in gamma-band power (40-80Hz) for V1 and V4. There were no statistically significant differences in power between groups. The shaded area around each curve indicates 95% confidence intervals.

### MEG and MRI Acquisition

MEG data were acquired using a 306-channel Neuromag MEG device (Vectorview, Elekta, Finland). A structural T1 brain scan was acquired for source reconstruction using a Siemens MAGNETOM Trio 3T scanner. MEG sensors were co-registered with anatomical MRI data by matching the digitised head-shape data with surface data from the structural scan (Jenkinson and Smith, 2001). For each participant, a cortical mesh was constructed using Freesurfer v5.3 (Fischl, 2012), and registered to a standard fs_LR mesh (Van Essen 2012). For more detailed instructions, see *Supplementary Information*.

### MEG Pre-Processing

MEG data were pre-processed using Maxfilter (tSSS, .9 correlation), which supresses external sources of noise (Taulu and Simola, 2006). Further pre-processing was performed in Matlab 2014b using the Fieldtrip toolbox v20161024 (Oostenveld *et al*., 2010). Data were band-pass filtered (0.5-250Hz, Butterworth filter) and band-stop filtered (49.5-50.5Hz; 99.5-100.5Hz) to remove power-line contamination and harmonics. Data were epoched into segments of 4s (1.5s pre, 1.5s post stimulus onset, with ±0.5s padding), demeaned and detrended. Trials containing artefacts (SQUID jumps, eye-blinks, head movement, muscle) were removed if the trial-by-channel magnetomer variance exceeded 8×10^−23^. This resulted in a group average of 60.2 trials for the ASD group and 61.9 trials for the control group. Four noisy MEG channels were removed from all analyses.

### Source-Level Power

Source analysis was conducted using a linearly constrained minimum variance beamformer (Van Veen *et al*., 1997), which applies a spatial filter to the MEG data at each vertex of the cortical mesh. Due to differences in noise between sensor-types, covariance matrix terms resulting from multiplying magnetomer and gradiometer data were removed. Beamformer weights were calculated by combining this covariance matrix with leadfield information, with data pooled across baseline and grating periods. Following tSSS, sensor-level data had a rank 64 or below, and therefore a regularisation parameter of lambda 5% was applied. Data were band-pass filtered between 40-80Hz (gamma) and 8-13Hz (alpha), and source analysis was performed separately. While gamma is typically defined as a wider range of frequencies, here we focussed on a 40-80Hz sub-range for an optimal signal-to-noise-ratio for source localisation. To capture induced rather than evoked visual power, a period of 0.3-1.5s following stimulus onset was compared with a 1.2s baseline period (1.5-0.3s before grating onset).

### ROI definition

To quantify directed connectivity within the visual system, we selected two regions of interest (ROI): visual area 1 (V1) and visual area 4 (V4), defined using HCP-MMP 1.0 atlas (Glasser *et al*., 2016) (Figure 1C). Both regions show stimulus-related changes in oscillatory power (Figure 1E-F) and demonstrate reliable patterns of hierarchical connectivity: V1-to V4 connectivity is feedforward; whereas V4-to-V1 connectivity is feedback (Bastos *et al*., 2015a, b; Michalareas *et al*., 2016). 12 vertices from posterior V1 were excluded to ensure clear anatomical separation of the ROIs. To obtain a single spatial filter for each ROI, we performed a principal components analysis on the concatenated filters encompassing V1 and V4, multiplied by the sensor-level covariance matrix, and extracted the first component (Schoffelen *et al*., 2017). Broadband (0.5-250Hz) sensor-level data was multiplied by this spatial filter to obtain “virtual electrodes”. Finally, the change in oscillatory power between grating and baseline periods was calculated using multi-tapers (Hoogenboom *et al*., 2006) from 1-140Hz, 0.5s time window, sliding in steps of 0.02s and ±8Hz frequency smoothing.

### V1-V4 Directed Connectivity

To quantify V1-V4 directed functional connectivity, we used a spectrally resolved non-parametric version of Granger Causality (GC) – a statistical technique which measures the extent to which one time series can predict another (Granger, 1969; Dhamala *et al*., 2008). Data from V1 and V4 (0.3-0.1.5s post-stimulus onset) were split into 0.4s epochs to enhance the accuracy of results, Fourier transformed (Hanning taper; 2Hz smoothing), and entered into a non-parametric spectral matrix factorisation procedure. GC was then estimated between 1-140Hz for each ROI pair and averaged across hemispheres. To create surrogate data (with no inter-regional connectivity), 0.4s-long time-series were produced with the same spectral properties as V1/V4, modelled using the first autoregressive coefficient (Colclough *et al*., 2015; see *Supplementary Information* for MATLAB code). GC was estimated between these surrogate V1-V4 time-series using the same procedure as for the actual data. GC spectra from the actual data were compared with surrogate GC spectra using cluster-based permutation tests (see *Statistical Analysis*).

Asymmetries in GC values were quantified using a Directed Asymmetry Index (DAI), originally defined in Bastos *et al*., (2015b), see below:

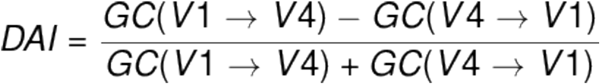

This results in normalised values (−1 to +1) for every frequency bin, with values above 0 indicating feedforward GC influence and values below 0 indicating feedback influence. DAI values were statistically compared between groups.

### Phase-Amplitude Coupling (PAC)

V1 time courses were examined for changes in alpha-gamma PAC. For detailed discussion about PAC computation and methodological issues see Seymour *et al*., (2017). Briefly, we calculated PAC between 7-13Hz phase (1Hz steps) and amplitudes 34-100Hz (in 2Hz steps), from 0.3-1.5s post-grating presentation. PAC values were corrected using 1.2 of data from the baseline period. In accordance with Seymour *et al*., (2017), we used a wide amplitude frequency range (34-100Hz) in order to characterise which gamma frequencies give rise to maximum changes in PAC (i.e. a data-driven approach). 34Hz was chosen as the lower limit of the range, as this is the lowest detectable amplitude frequency for phases from 7-13Hz. Amplitude-phase comodulograms (size: 33*7), were statistically compared between groups using cluster-based permutation testing (Maris and Oostenveld, 2007).

PAC was calculated using two separate approaches, a mean vector length algorithm (Özkurt and Schnitzler, 2011), MVL-Ozkurt, and a phase-locking algorithm (Cohen, 2008), PLV-Cohen. This decision was based on our previous study which compared the efficacy of four different PAC algorithms for MEG data analysis (Seymour *et al*., 2017). Additional details are outlined in the *Supplementary Information*, and code used for PAC computation is available at: https://github.com/neurofractal/sensory_PAC.

### Statistical Analysis

Statistical analysis was performed using cluster-based permutation tests (Maris and Oostenveld, 2007), which consist of two parts: first an independent-samples t-test is performed, and values exceeding an uncorrected 5% significance threshold are grouped into clusters. The maximum t-value within each cluster is carried forward. Second, a null distribution is obtained by randomising the condition label (e.g. ASD/control) 1000 times and calculating the largest cluster-level t-value for each permutation. The maximum t-value within each original cluster is then compared against this null distribution, and the null hypothesis is rejected if the test statistic exceeds a threshold of p<.05. Cluster-based permutation tests are an effective way to address the multiple-comparison problem for neuroimaging data, which is especially problematic for M/EEG data analysed over frequency, time *and* space (Maris and Oostenveld, 2007).

## Results

### Oscillatory Power

The change in oscillatory power following presentation of the visual grating, versus baseline, was calculated on a cortical mesh for the alpha (8-13Hz) and gamma (40-80Hz) bands. For both ASD and control groups there was a statistically significant relative increase in gamma power (Figure 1B) and a relative decrease in alpha power (Figure 1C), localised to the ventral occipital cortex. This replicates previous MEG/EEG studies using visual grating stimuli (Hoogenboom *et al*., 2006; Michalareas *et al*., 2016). Interestingly, there were no significant differences in relative gamma or alpha power between groups (p>.05, see Supplementary Figure S1).

Two regions of interest (ROI) were defined in V1 and V4 (Figure 1D). Changes in oscillatory power (grating vs baseline) from V1 (Figure 1E) and V4 (Figure 1F) showed characteristic increases in gamma-band power (40-80Hz) and decreases in alpha/beta power (8-20Hz). Between groups, there were minor differences between the power spectra, including a larger alpha/beta induced power change for the ASD group (Fig 1E, 1F, purple line) but none of these differences were significant (both p>.05).

In sum, we found no evidence for group differences (control vs ASD) in gamma or alpha relative oscillatory power following the presentation of a visual grating. Additionally, there were no significant correlations between oscillatory power in V1/V4 and behavioural Autism Quotient (AQ) scores for the ASD group (see Supplementary Figure S2).

### Feedforward / Feedback Connectivity

The directed functional connectivity between V1-V4 was quantified using Granger Causality (GC). Across groups, all reported increases in bidirectional V1-V4 GC were greater than for surrogate data (Supplementary Figure S3). For the control group (Figure 2A), V1-to-V4 (henceforth termed feedforward) connectivity showed a prominent increase from 40-80Hz in the gamma band. In contrast, V4-to-V1 (henceforth termed feedback) connectivity showed a prominent increase from 8-13Hz in the alpha band (Figure 2A). This dissociation between feedforward gamma and feedback alpha, replicates previous findings in macaques and humans (Bastos *et al*., 2015b; Michalareas *et al*., 2016). The feedforward gamma-band peak (40-80Hz) was also evident in the ASD Granger spectra (Figure 2B, red line). There was a reduction in the alpha-band feedback peak in the ASD group compared with controls (Figure 2B, blue line).

**Figure 2:**
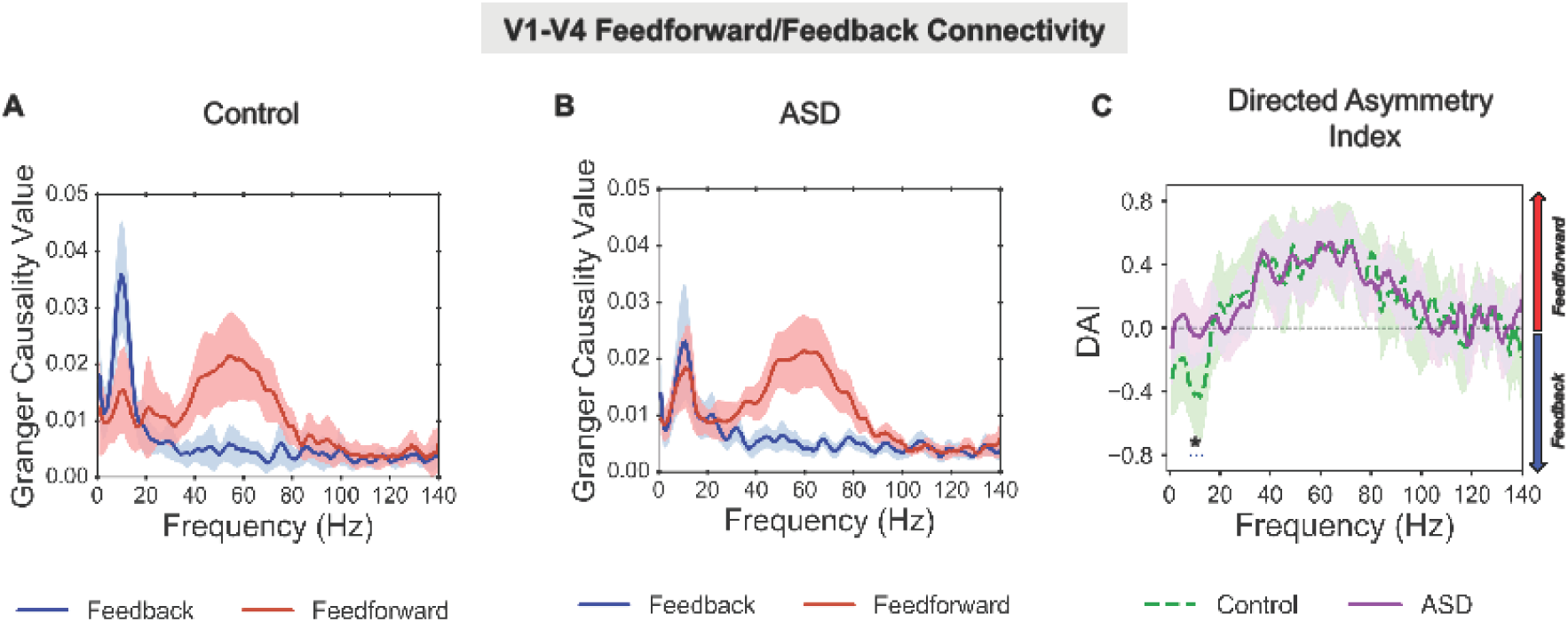
V1-V4 Feedforward/Feedback Connectivity. (A) For the control group there was a peak in granger causality (GC) values, in the gamma-band (40-80Hz, red line) for V1-to-V4 feedforward connectivity, and a peak in GC values in the alpha band (8-13Hz, blue line) for V4-to-V1 feedback connectivity. (B) For the ASD group there was also a peak in GC values in the gamma-band for V1-to-V4 feedforward connectivity, however there was a smaller peak in GC in the alpha-band for V4-to-V1 feedback connectivity. For comparisons with surrogate data per group, please see Supplementary Figure S3. (C) The difference between feedforward and feedback connectivity was quantified as the directed asymmetry index (DAI, see Material and Methods). The difference in DAI between control (dashed, green line) and ASD (solid, purple line) was significant (p=.036), with lower DAI values (p=.036) between 8-14Hz for the control group, suggesting reduced V4-to-V1 feedback connectivity in autism. The shaded area around each GC line indicates 95% confidence intervals.

To quantify asymmetries in feedforward/feedback connectivity between groups, we calculated the directed asymmetry index (DAI, see *Materials and Methods*). The control group displayed a feedback peak from 0-20Hz (negative DAI values) and feedforward peak from 40-80Hz (positive DAI values). By statistically comparing DAI between groups, it was found that values from 8-14Hz were significantly lower (p=.032) for the control group than the ASD group. All other frequencies, including gamma (40-80Hz) showed similar DAI values between groups. This suggests reduced V4-to-V1 feedback connectivity for the ASD group, mediated by alpha-band oscillations (8-14Hz), but typical V1-to-V4 feedforward connectivity mediated by gamma oscillations (40-80Hz).

There was no feedforward Granger causality peak in the theta-band (4-8Hz) for either the control or ASD group, as previously reported using ECoG (Spyropoulos *et al*., 2018). This could be due to lower sensitivity of MEG recordings (Michalareas *et al*., 2016), as well as the centrally-masked visual grating (Fig. 1A).

### Alpha-Gamma Phase Amplitude (PAC) in V1

Activity from visual area V1 was examined for changes in alpha-gamma PAC using two separate approaches (MVL-Ozkurt; PLV-Cohen, see Materials and Methods). Frequency comodulograms showed increased PAC in the control group, peaking at 8-10Hz phase frequencies and 50-70Hz amplitude frequencies (Figure 3A,B). These results replicate Seymour *et al*., (2017), who showed increased alpha-gamma PAC in an adult population using the same visual grating stimulus. The comodulograms for the ASD group displayed lower PAC values, with no clear positive peak (Figure 3C,D). Comparing control vs. ASD groups, there was a single positive cluster of greater PAC between 8-9Hz and 52-74Hz for the MVL-Özkurt approach (Figure 3E, p=.029); and a single positive cluster between 8-9Hz and 54-74Hz for the PLV-Cohen approach (Figure 3F, p=.037). This suggests that the coupling between alpha and gamma oscillations during perception in primary visual cortex is reduced in autism. The similarity in PAC comodulograms between MVL-Özkurt and PLV-Cohen approaches, indicates that the results generalise across both PAC metrics.

**Figure 3:**
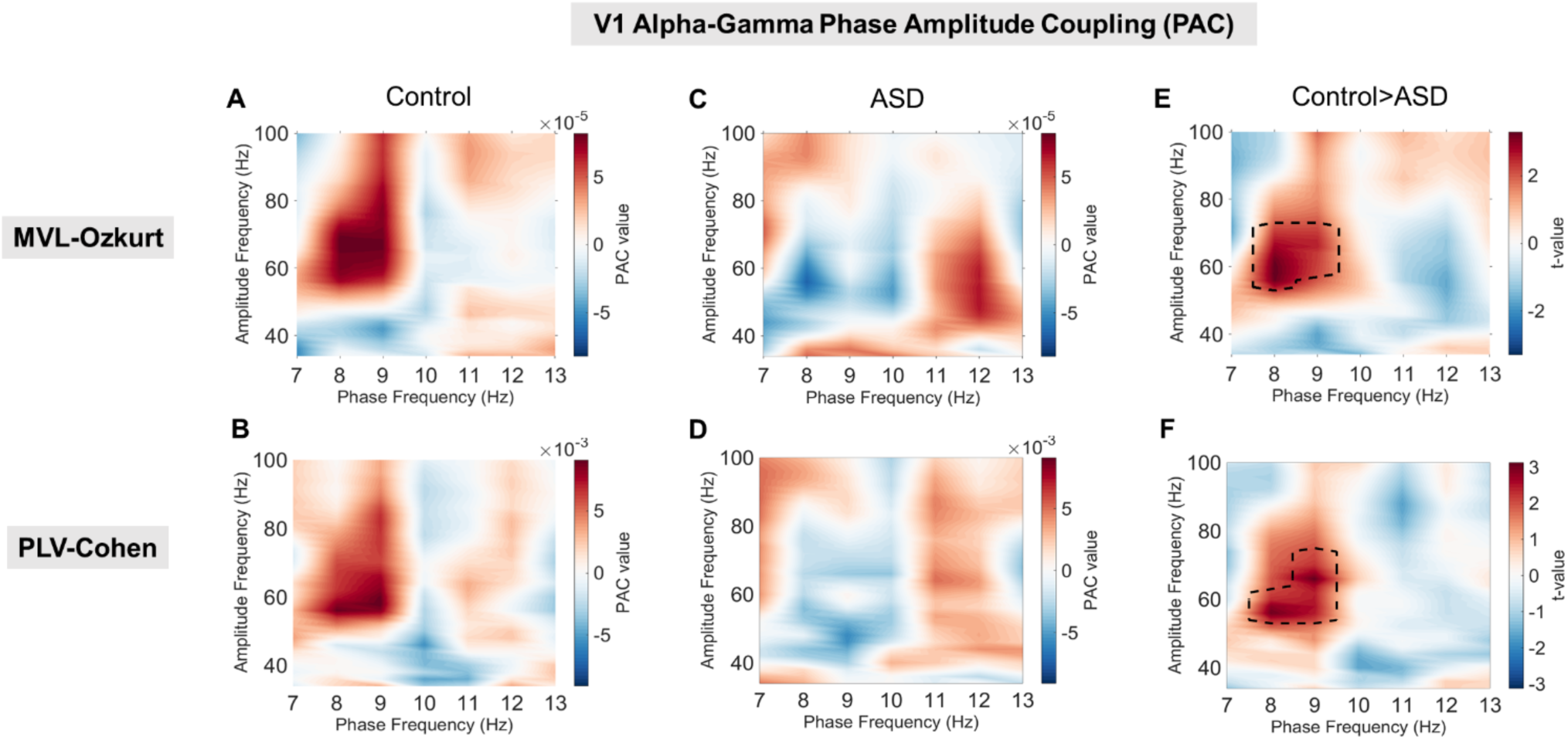
V1 Phase Amplitude Coupling using the MVL-Özkurt (A, C, E) and the PLV-Cohen (B, D, F) approaches (see Materials and Methods). (A, B) The control group showed increased alpha-gamma PAC compared with baseline, with a peak between 50-80Hz amplitude and 7-9Hz phase. (C, D) The ASD group showed less prominent increases in PAC with a much smaller peak from 40-70Hz amplitude and 11-13Hz phase shown in C and an even smaller peak shown in D. (E, F) Robust statistical comparison (see Materials and Methods for details) indicated significantly larger PAC for the control compared to the ASD group (p=.029 in E and p=.037 in F) from 54-72Hz amplitude and 8-9Hz phase.

### Connectivity - Behaviour Correlation

Behavioural ASD data from the Autism Quotient (AQ) and Glasgow Sensory Questionnaires (GSQ; Baron-Cohen *et al*., 2001a; Robertson and Simmons, 2013) were correlated with group differences in alpha-band DAI and alpha-gamma PAC (Figure 5). The AQ questionnaire measures general autistic traits, whilst the GSQ measures the level of reported sensory hypo- and hyper-sensitives across domains (Baron-Cohen *et al*., 2001a; Robertson and Simmons, 2013). There was a significant positive correlation between AQ score and alpha DAI (Figure 4B, r=.526, p=.025) suggesting that increased V4-to-V1 feedback connectivity (negative DAI values) is related to lower levels of autistic traits (lower AQ scores). There were no other significant correlations for the GSQ or PAC.

**Figure 4:**
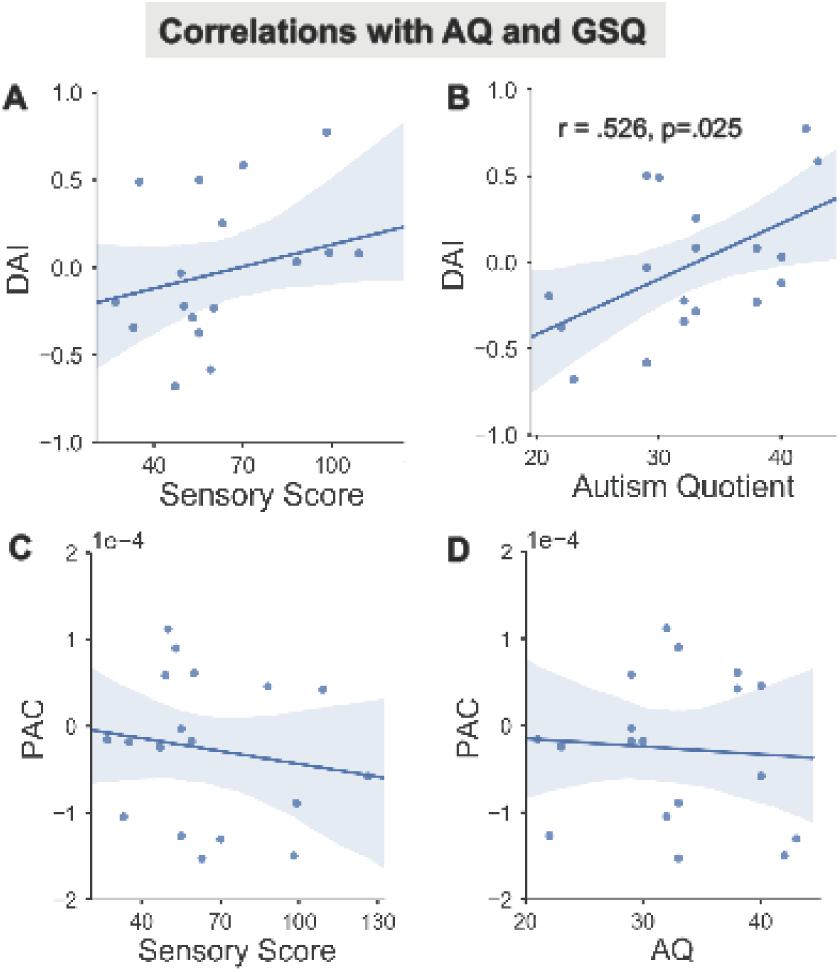
For the ASD group, the correlation between alpha-band DAI (A-B), alpha-gamma PAC (C-D) and Autism Quotient (B,D), Glasgow Sensory Score (A,C) was plotted with regression line (95% confidence interval indicated by shaded region). (B) There was a positive correlation between DAI and AQ score.

This analysis was repeated for the behavioural data from the control group. However, there were no significant correlations for any combination of DAI/PAC and AQ/GSQ data (see Supplementary Figure S5), p<.05.

## Discussion

This study examined the oscillation-based functional connectivity within the visual system of autistic adolescents and typically developing age-matched controls. Confirming our hypotheses (Kessler *et al*., 2016), we found a reduction in alpha-band (8-13Hz) feedback connectivity from V4-to-V1 in the ASD group alongside a reduction in the coupling between alpha and gamma oscillations in V1, measured via PAC, suggesting dysregulation of local connectivity in autism. Further in agreement with predictions (Kessler et al., 2016) aberrant connectivity patterns were observed in the absence of significant group differences in oscillatory power-changes relative to baseline (Figure 1; Supplementary Figure S1).

### Feedback / Feedforward Connectivity

By examining frequency-specific asymmetries in V1-V4 connectivity during visual processing (Bastos *et al*., 2015; Michalareas *et al*., 2016), this study found that the ASD group had specific reductions in feedback, but not feedforward, connectivity. This is consistent with previous MEG and fMRI studies showing a reduction in global connectivity in autism (Hughes, 2007; Khan *et al*., 2013; Kitzbichler *et al*., 2015). Having said this, it should be acknowledged that connectivity between visual regions V1-V4 might be better characterised as “inter-regional” rather than truly global. Future ASD-MEG research could examine global feedback/feedforward connectivity using measures of directed functional connectivity (e.g. Granger Causality) in concert with higher-level cognitive tasks involving a more extended set of cortical regions.

Using a simple visual paradigm, this study did not reveal an increase in connectivity from V1-to-V4 for the ASD group mediated by gamma oscillations, suggesting equivalent levels of feedforward information flow in the visual system between groups. Whilst Khan and colleagues reported increased feedforward connectivity in autism (Khan *et al*., 2015), they focussed on somatosensory rather than visual processing with a younger group of adolescent participants. In any case, we hypothesise that where visual processing can be achieved via feedforward processes (reflected at gamma frequencies), autistic participants may perform on par or even outperform their typically developing peers (Mottron *et al*., 2006). For example, during visual search tasks autistic participants have been reported to perform faster than controls (Jobs *et al*., 2018).

In contrast, we observed a reduction in feedback connectivity from V4 to V1 that was specific to alpha-band oscillations (8-14Hz, see Figure 2). While a comparison with surrogate data (Supplementary Figure S3) revealed a significant alpha feedback peak for the ASD group, it did not differ from the alpha feedforward peak, resulting in a DAI significantly closer to 0 than in the control group (Fig. 2C). Our data suggest that whilst relative alpha power was unaffected (Fig. 1 C, E, F; Supplementary Figure S1), the feedback flow of information from higher to lower visual regions was reduced in our ASD sample. A reduced ability to implement top-down modulation of bottom-up visual information may result in the atypical visual processes reported by many autistic individuals (for reviews see, Kessler *et al*., 2016; Simon and Wallace, 2016).

Despite observing no significant correlation between oscillatory power and AQ or GSQ at any frequency (see Supplementary Figure S2), a significant correlation was revealed between the reduction in alpha feedback connectivity and AQ score in the ASD group, further supporting our hypothesis of decreased top-down connectivity in ASD. However, we did not find a corresponding correlation with GSQ score that would corroborate our hypothesis with respect to the severity of sensory symptoms. A possible reason for the lack of correlation could be that the GSQ is a general questionnaire, which addresses aberrations across seven sensory domains (visual, auditory, gustatory, olfactory, tactile, vestibular, proprioceptive) at the expense of an in-depth assessment of any specific modality. In addition, different items per domain address either hypo- or hyper-sensitivities (resulting in only 3 items per expression, per domain) and the obtained scores in our sample indeed reflect a mix between both symptom expressions (see Supplementary Figure S6). This and the observation that sensory symptoms were only reported as “rarely” or “sometimes” in our sample, may have added to a variable relationship between brain measures and GSQ scores. In conclusion, brain-behaviour relationships might be better assessed using more precise psychophysical tests of visual perception (Ashwin et al., 2009), combined with formal clinical assessments.

### PAC

Within primary visual cortex (V1), there was a reduction in alpha-gamma PAC for the ASD group (Figure 3). It is important to note that the group differences in PAC arose despite similar relative changes in gamma and alpha power (Figure 1). Interestingly, one previous ASD study reported reduced inter-regional connectivity and local alpha-gamma PAC during face processing, despite similar event-related activity and oscillatory power between groups (Khan *et al*., 2013). As reviewed in the Introduction, reports of gamma band responses (GBA) in ASD are inconsistent, with some M/EEG studies reporting hyper-reactivity (Orekhova *et al*, 2007; Cornew *et al*, 2012), while others reporting reduced GBA at the local level (e.g. Khan *et al*, 2013). Future studies should therefore explore the precise regulation of gamma oscillations via cross-frequency coupling, rather than relying on measures of power alone (Canolty and Knight, 2010; Kessler *et al*., 2016; Simon and Wallace, 2016).

PAC has indeed been reported to rely heavily on local inhibitory populations of neurons (Onslow et al., 2014) and could therefore be a more reliable indicator of an E-I imbalance in ASD than GBA. The observed reduction in PAC is therefore consistent with histological findings showing underdeveloped inhibitory interneurons (Casanova et al., 2003) and an E-I imbalance in autism (Rubenstein and Merzenich, 2003). Affected local inhibitory processes could manifest as high-frequency ‘noisy’ activity and reduced signal-to-noise in perceptual systems, as reported in ASD (Casanova *et al*., 2003; Rubenstein and Merzenich, 2003; Vilidaite *et al*., 2017). However, it should be noted that further corroborating evidence will be required before a definitive link between PAC and E-I interactions can be established.

It has been proposed that dysregulated local activity could have concomitant effects on establishing patterns of inter-regional and global connectivity (Voytek and Knight, 2015). In the context of our current investigation of the autistic visual system, reduced local PAC in V1 could therefore reveal a dysfunctional relationship with V1-V4 interregional connectivity. Indeed, an exploratory analysis reported in Supplementary Figure S4 revealed a correlation between negativity of DAI (=predominance of alpha feedback connectivity) and the strength of PAC across groups. Whilst the control group in its majority showed increased feedback alpha and increased alpha-gamma PAC, the relationship for the ASD group was significantly more variable (Supplementary Figure S4). However, due to the employed visual grating paradigm and the limited samples tested here, future research is required to test the general claims that PAC acts as a general cortical mechanism for oscillatory multiplexing to link connectivity at the global and local scales (Canolty and Knight, 2010; Seymour *et al*., 2017) and that this mechanism is specifically affected in autism (Kessler *et al*., 2016).

Interestingly, we did not find a relationship between AQ or GSQ and PAC in the ASD group (Figure 4 C, D), although there was a relationship with alpha DAI (Supplementary Figure S4). In addition to the discussed issues regarding sensitivity of the GSQ, PAC may be related to specific clinical features of autism rather than general autistic traits (see *Limitations*). Accordingly, a recent study reported a correlation between the social component of the Autism Diagnostic Observation Schedule (ADOS) and local PAC in an adolescent autistic sample (Mamashli *et al*., 2018).

### Neurocognitive Models of Perception

Our results link with emerging theories of typical perception. Predictive-coding accounts of cortical activity describe the passage of top-down predictions from higher to lower areas via feedback pathways, with prediction errors computed at each level of the hierarchy being passed forward via feedforward pathways (Friston, 2005). Predictive-coding accounts of autism suggest that differences in perception emerge from fewer or hyper-precise top-down predictions, such that perception is less influenced by prior knowledge and contextual cues (Pellicano and Burr, 2012; Palmer *et al*., 2017). Despite limitations, our data support this proposal by showing reduced feedback connectivity in the visual cortex in autism. We propose that where top-down information flow is reduced, the perceptual system could be forced from predictive to reactive, with increased prediction error signalling and concomitant impacts on autistic symptoms (Kessler *et al*., 2016). This is supported by the observed correlations between feedback connectivity (DAI) and AQ score (Figure 4B) and between DAI and PAC (Supplementary Figure S4) but requires thorough further investigation.

### Clinical Implications and Limitations

We note three limitations to this study. First, we did not collect a formal clinical assessment of autism, e.g. the ADOS. We therefore implemented strict participant exclusion criteria, only including autistic participants with a confirmed clinical diagnosis of ASD or Asperger’s syndrome. Between groups, there were significant differences in autistic and sensory traits (Table 1). However, upon closer inspection of GSQ data, the ASD group showed a mixture of hyper- and hypo-sensitive traits between different sensory modalities making precise brain-behavioural correlations problematic (Supplementary Figure S6). This may explain the lack of relationship between oscillatory connectivity and GSQ scores in autism (Figure 5A, C). Brain-behaviour relationships might be better assessed using psychophysical tests of visual perception (Ashwin *et al*., 2009), combined with formal clinical assessments. Second, due to the relatively low number of participants tested in each group, it would be inappropriate to generalise our findings, at this time, to the entire ASD spectrum and beyond the current visual grating paradigm. In addition, a greater number of participants may be required to achieve the appropriate statistical power for brain-behaviour correlations. Nonetheless, our novel analysis approach has revealed interesting and predicted findings (Kessler *et al*., 2016) despite a quite diverse high-functioning ASD sample (e.g. GSQ scores) and may therefore provide important findings, upon which future research can replicate and extend. Third, we constrained our connectivity analyses to two regions of interest (V1, V4) located early in the visual system, due to their hierarchical connectivity, and the low-level nature of the visual grating stimulus. However, we may have missed the opportunity to characterise more complex feedforward-feedback relationships in wider visual cortex. Future work should therefore include more ROIs in combination with stimuli requiring participants to explicitly engage in feedback processing to constrain visual perception. This approach could be particularly useful with high-functioning individuals, and help characterise the neurophysiological basis of autistic perception (Kessler *et al*., 2016; Robertson and Baron-Cohen, 2017).

## Abbreviations

ASD: Autism Spectrum Disorder
MEG: Magnetoencephalography
PAC: Phase Amplitude Coupling
E-I: Excitation-Inhibition
ECoG: Electrocorticography
GSQ: Glasgow Sensory Questionnaire
AQ: Autism Quotient
GC: Granger Causality
DAI: Directed Asymmetry Index
ROI: Region of Interest
tSSS: temporo-spatial signal separation

## Data availability

The data that support the findings of this study are available on reasonable request from corresponding author, RS, in a pre-processed and deanonymized form. The raw data are not publicly available due to ethical restrictions. MATLAB data analysis code for this study will be made available openly on Github after manuscript acceptance. Code for PAC computation is openly available at: https://github.com/neurofractal/sensory_PAC.

## Acknowledgments

We wish to thank: the volunteers who gave their time to participate in this study; Dr. Jon Brock for intellectual contributions to experimental design; and the Wellcome Trust, Dr Hadwen Trust and Tommy’s Fund for supporting research costs. Robert Seymour was supported by a cotutelle PhD studentship from Aston University and Macquarie University. Jan-Mathijs Schoffelen was supported by The Netherlands Organisation for Scientific Research (NWO Vidi: 864.14.011).

## Competing interests

The authors have no competing interests to disclose. A version of this manuscript has been uploaded to the pre-print BioRxiv service.

## Supplementary Information

### Supplementary Methods

#### Participant Exclusion

MEG data from a further 9 participants was collected but excluded, due to: intolerance to MEG (2 ASD); movement over 0.5cm (2 ASD, 2 control); metal artefacts (1 ASD, 1 control); AQ score over 30 (1 control).

#### MEG Acquisition

MEG data were acquired using a 306-channel Neuromag MEG system (Vectorview, Elekta, Finland) made up of 102 triplets of two orthogonal planar gradiometers and one magnetometer. All recordings were performed inside a magnetically shielded room at a sampling rate of 1000Hz. Five head position indicator (HPI) coils were applied for continuous head position tracking, and visualised post-acquisition using an in-house Matlab script. For MEG-MRI coregistration purposes three fiducial points, the locations of the HPI coils and 300-500 points from the head surface were acquired using the integrated Polhemus Fastrak digitizer. Visual stimuli were presented on a screen located 86cm from participants (resulting in 2 cycles/degree for the visual grating), and auditory feedback through MEG-compatible earphones.

#### Structural MRI

A structural T1 brain scan was acquired for source reconstruction using a Siemens MAGNETOM Trio 3T scanner with a 32-channel head coil (TE=2.18ms, TR=2300ms, TI=1100ms, flip angle=9°, 192 or 208 slices depending on head size, voxel-size = 0.8×0.8×0.8cm).

#### MEG-MRI Coregistration and 2D Cortical Mesh Construction

MEG data were co-registered with participants MRI structural scan by matching the digitised head-shape data with surface data from the structural scan (Jenkinson and Smith, 2001). Two control participants did not complete a T1 structural MRI and therefore a pseudo-MRI was used, see Gohel *et al*., (2017) for full procedure. The aligned MRI-MEG images were used to create a forward model based on a single-shell description of the inner surface of the skull (Nolte, 2003), using the segmentation function in SPM8 (Litvak *et al*., 2011). The cortical mantle was then extracted to create a cortical mesh, using Freesurfer v5.3 (Fischl, 2012), and registered to a standard fs_LR mesh, based on the Conte69 brain (Van Essen 2012), using an interpolation algorithm from the Human Connectome Project (Van Essen et al., 2012; instructions here: https://goo.gl/3HYA3L). Finally, the mesh was downsampled to 4002 vertices per hemisphere.

#### PAC

The mean vector length approach estimates PAC from a signal with length N, by combining phase (ϕ) and amplitude information to create a complex-valued signal: 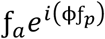 (Canolty *et al*., 2006), in which each vector corresponds to a certain time-point (n). If the resulting probability distribution function is non-uniform, this suggests a coupling between ƒ_p_ and ƒ_a_, which can be quantified by taking the length of the average vector. As recommended by Özkurt & Schnitzler (2011) a normalisation factor was also applied corresponding to the power of ƒ_a_.

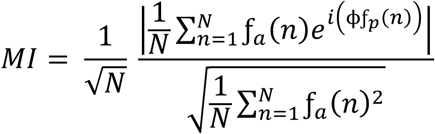

The PLV-Cohen approach assumes that if PAC is present, the envelope of ƒ_a_ should oscillate at the frequency corresponding to ƒ_p_. The phase of ƒ_a_ envelope can be obtained by applying the Hilbert transform (angle): ϕƒ_a_. The coupling between the low-frequency ϕƒ_p_ phase values and the phase of the amplitude envelope, ϕƒ_a_, can be quantified by calculating a phase locking value (PLV), in much the same way as determining phase synchronisation between electrophysiological signals.

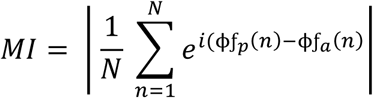

#### Surrogate Data

To create surrogate data we used the MATLAB code shown below. Functions are based on MEG-ROI Nets repository (Colclough et al., 2015) which can be found at: https://github.com/OHBA-analysis/MEG-ROI-nets/tree/master/%2BROInets.

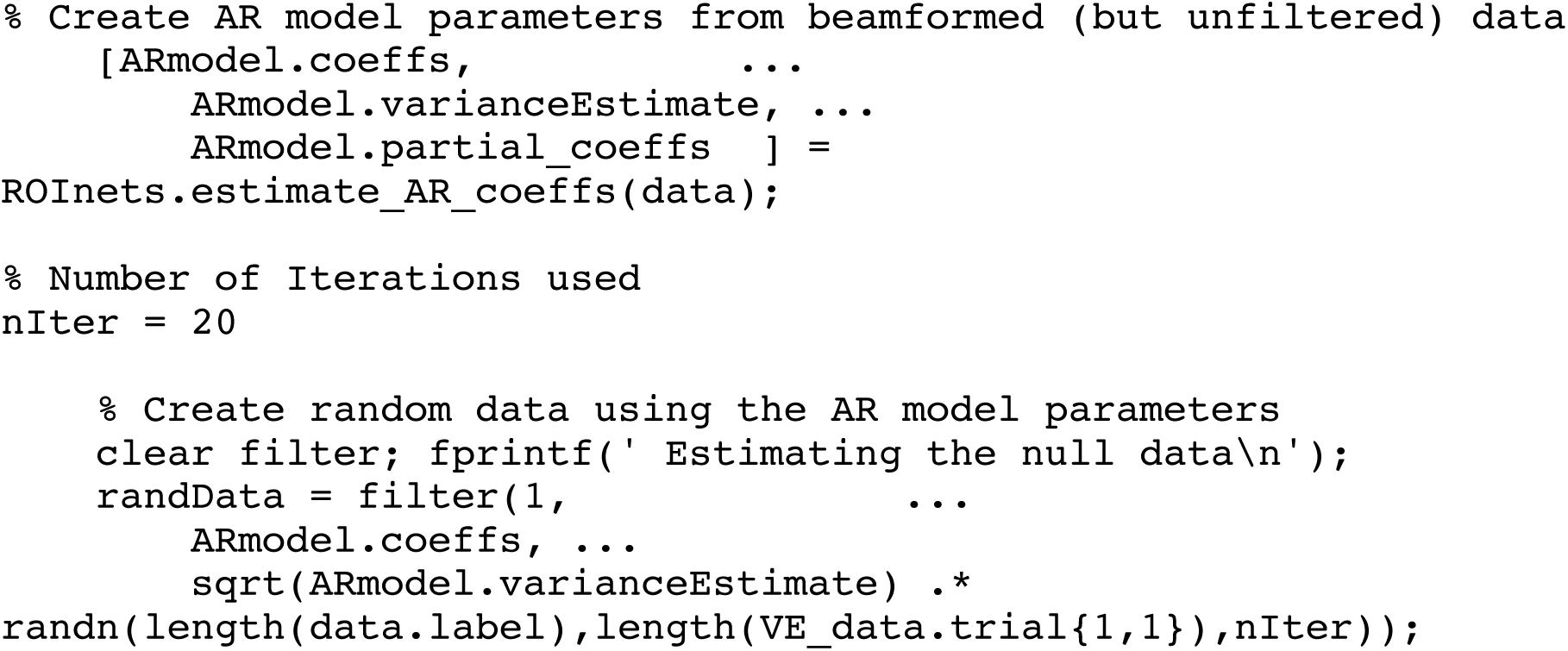

## Supplementary Figures

**Figure S1:**
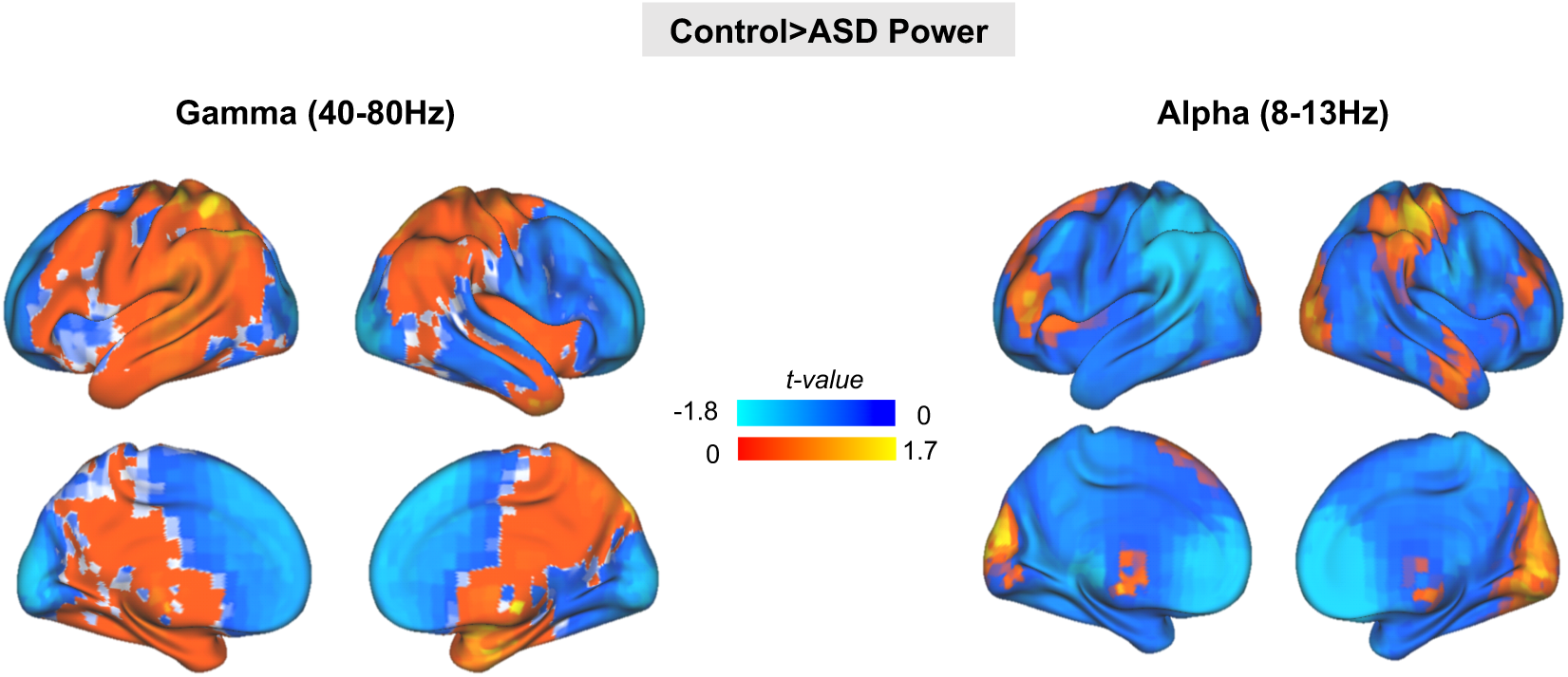
Brain-wide statistical comparison of control>ASD source-space oscillatory power for gamma (40-80Hz) and alpha (8-13Hz). There were no significant differences in either alpha or gamma power between groups (p>.05, corrected for multiple comparisons).

**Figure S2:**
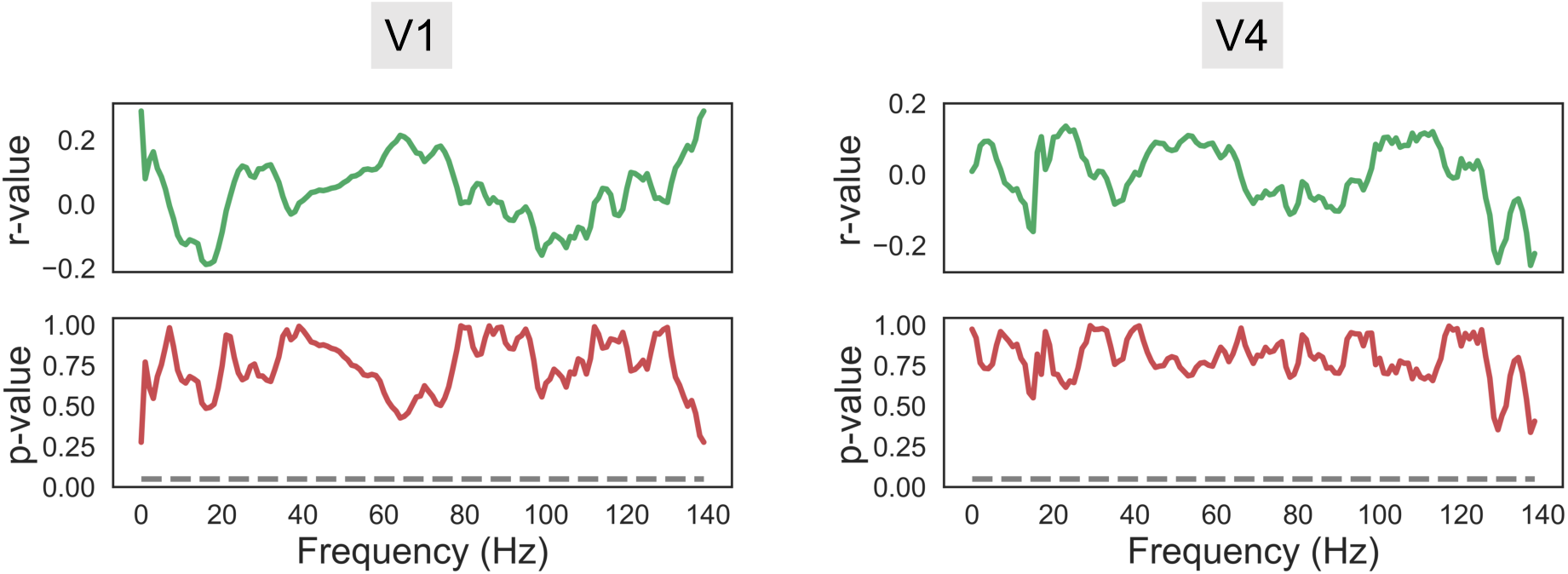
The correlation between V1 and V4 power spectrums (change in power between baseline and grating time-periods) and Autism Quotient (AQ) scores was calculated for the ASD group using Pearson’s r. No frequency passed an uncorrected p<.05 significance threshold (indicated by the grey dotted line).

**Figure S3:**
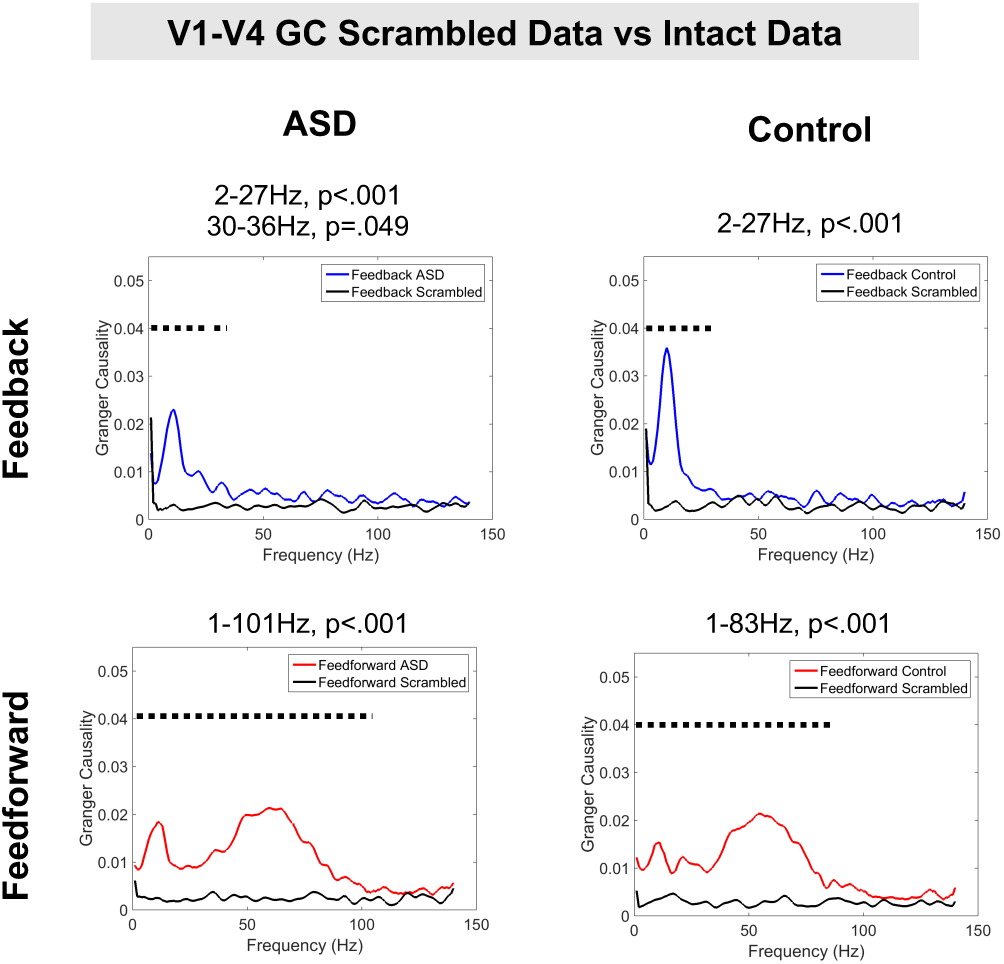
V1-V4 feedforward and V4-V1 Granger Causality (GC) values were statistically compared with GC values computed using scrambled V1/V4 data with same spectral properties as the intact data. On each sub-figure the black dotted line signifies intact GC values significantly greater than scrambled GC values (p<.05). The exact frequency range and p-values are listed at the top of each plot.

**Figure S4:**
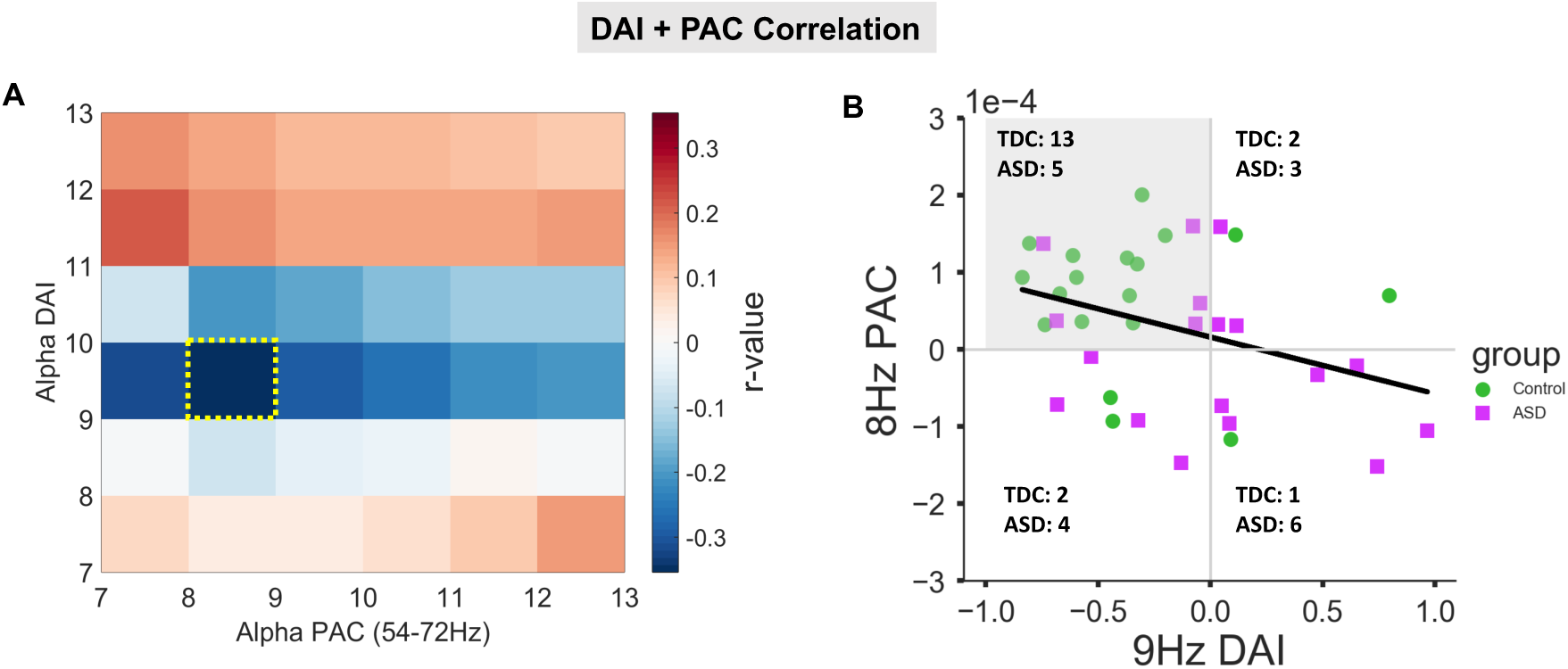
(A) To investigate the correlation between feedback connectivity and PAC, a cross correlation matrix was calculated pooled across ASD and control participants in 1Hz steps between alpha PAC, averaged between 54-72Hz, and 7-13Hz directed asymmetry index (DAI). This produced a negative correlation peak, shown with yellow box, at 8Hz PAC, 9Hz DAI. (B) The correlation between 8Hz PAC, 9Hz DAI is negative across both groups (Pearson’s r = -.35, p = .034). Note that only the top-left quadrant reflects the relationship postulated for effective processing: feedback alpha in form of negative DAI values (x-axis) paired up with positive PAC values (y-axis). 13 out of 18 control participants but only 5 out of 18 ASD participants are located in this quadrant (frequencies per group are indicated for each quadrant). The frequency distribution across the four quadrants was significantly different between the two groups as revealed by a Chi-Square test (chisq=7.99; p=.046), differing most strongly in the top-left (control: 13; ASD: 5) and bottom-right (control: 1; ASD: 6) quadrants.

**Figure S5:**
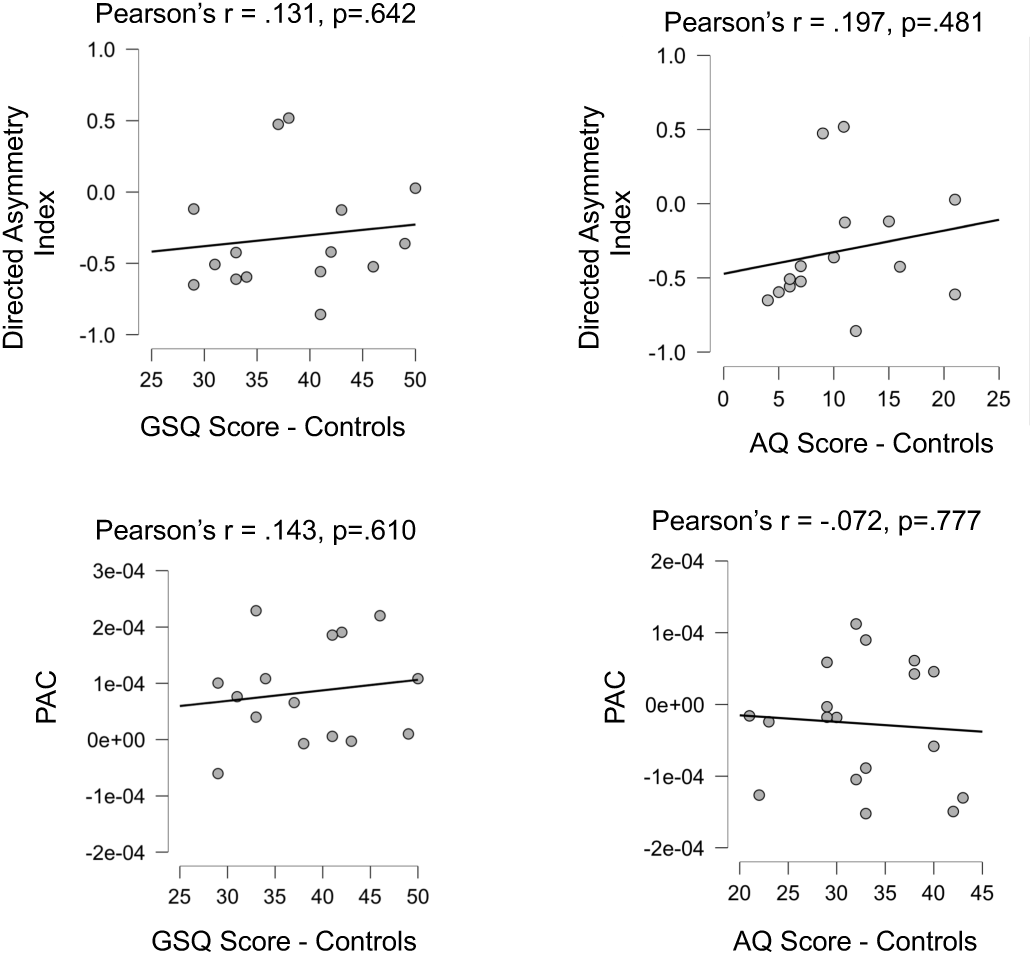
The correlation between Autism Quotient (AQ) score for the control group and the Directed Asymmetry Index (DAI) was calculated using JASP. For the control group, the correlation between alpha-band DAI, alpha-gamma PAC and Autism Quotient, Glasgow Sensory Score was plotted with regression line. There were no significant correlations, p>.05.

**Figure S6:**
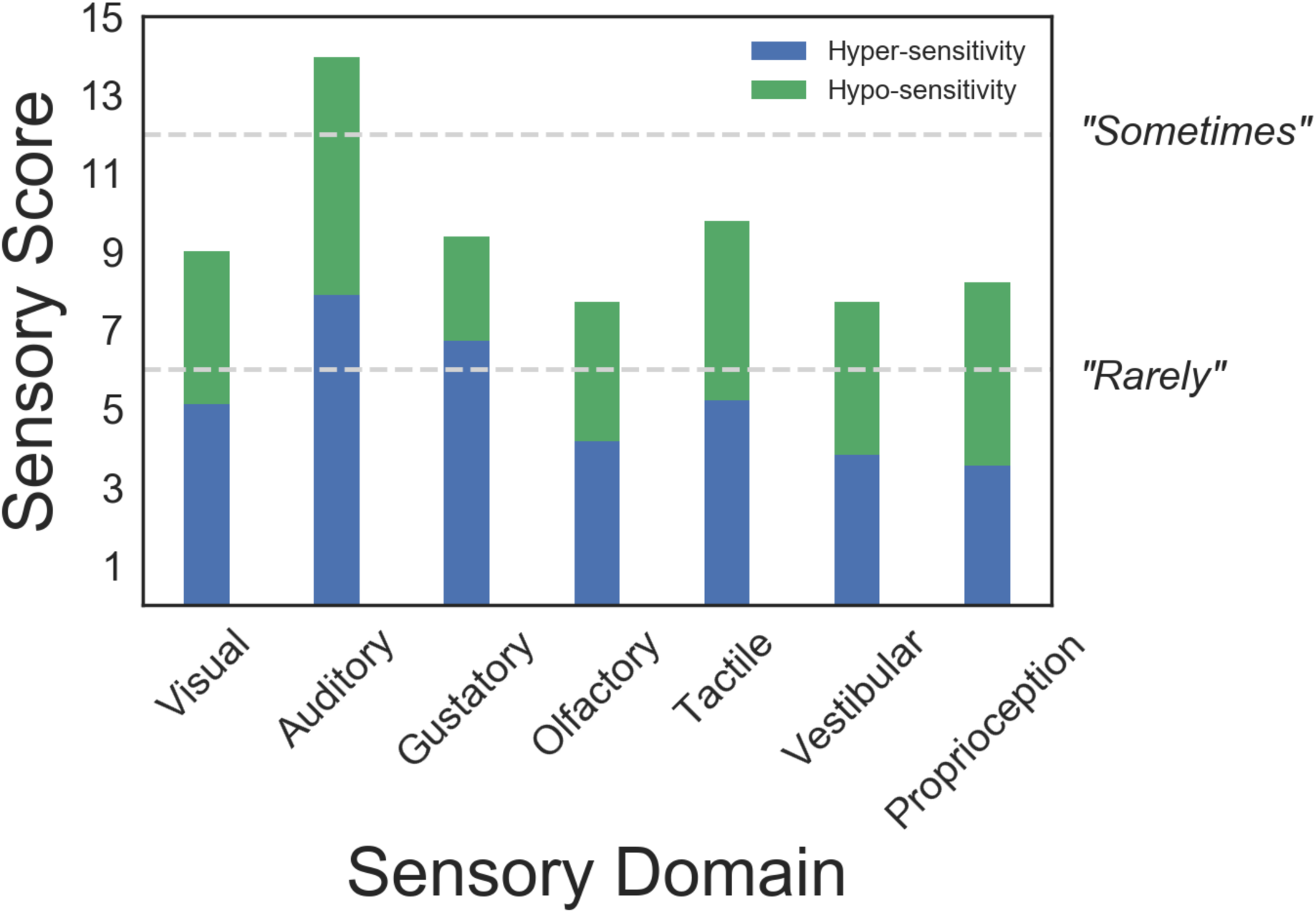
Responses to the Glasgow Sensory Questionnaire were grouped by sensory domain (maximum score = 20) and hypo- / hyper-sensitivity (green and blue bars respectively). Our data show a heterogeneous pattern of sensory symptoms, with mixture of hypo- and hyper-sensitivities. Visual symptoms scored 9.0/20 corresponding to questionnaire answers between “Rarely” and “Sometimes”. Auditory sensory symptoms were higher than for other modalities.

## Notes

#### Summary of Updates

Responded to reviewers comments about interpretation of data as presented in discussion section

